# Nano-org, a functional resource for single-molecule localisation microscopy data

**DOI:** 10.1101/2024.08.06.606779

**Authors:** S. Shirgill, D.J. Nieves, J.A. Pike, M.A. Ahmed, M.H.H. Baragilly, K. Savoye, J. Worboys, K.S. Hazime, A. Garcia, D.J. Williamson, P. Rubin-Delanchy, R. Peters, D.M. Davis, R. Henriques, S.F. Lee, D.M. Owen

## Abstract

We present a publicly accessible, curated, and functional resource, termed “nano-org”, containing single-molecule localisation microscopy (SMLM) data representing the nanoscale distributions of proteins in cells. Nano-org is searchable by comparing the statistical similarity of the datasets it contains. This unique functionality allows the resource to be used to understand the relationships of nanoscale architectures between proteins, cell types or conditions, enabling a new field of spatial nano-omics.

---

The nanoscale organisation and oligomerisation of proteins are critical processes in fundamental biology. For instance, nanoscale protein clustering is a key mechanism in regulating signal transduction by orchestrating protein-protein interaction rates [1]. Moreover, the organisation of cytoskeletal components, such as the nanoscale architecture of cortical actin, helps define cell mechanical properties [2,3]. Aberrant protein nanoscale organisation has been implicated in diseases, for example Alzheimer’s disease and type II diabetes [4]. Given this importance, there is a need for a resource to allow researchers to compare protein nanoscale organisations. High quality community-driven accessible databases/atlases have been transformative across biology in the areas of predictive protein structure, cell phenotyping and large-scale omics, such as NCBI and PDB [5, 6]. However, these platforms have yet to be utilised for comparing protein distributions and assemblies - a field termed spatial nano-omics. Single-molecule localisation microscopy (SMLM) is a fluorescence microscopy technique that provides coordinates of protein distributions in cells with nanometre precision [7]. While SMLM databases exist, they lack two features required to enable spatial nano-omics: curation and functionality [8]; features that are required to turn a repository into a resource. To address this need, we present nano-org, a publicly accessible and curated database for SMLM data.

Nano-org is freely accessible at nano-org.bham.ac.uk. Uploading and downloading data requires registration and email verification. When processing files containing the coordinates of SMLM localisations it uses standardised tiled 3 × 3μ m regions-of-interest (ROIs), facilitating the application of the similarity algorithm implemented in the resource. These can be uploaded manually or produced automatically from full field-of-view (FOV) data. Optionally cell bounding polygons can be uploaded alongside full FOV data. Datasets are organised into folders representing experimental conditions and are accompanied by metadata such as the SMLM modality, protein identity, cell type, fluorophore tag and other relevant information, including DOI link for published data. The metadata also includes whether the data has undergone drift and multiple-blink correction.

To ensure the validity of downstream data analysis, data stored on nano-org is curated and subject to restriction. It currently accepts 2D datasets in .csv format acquired from one of three SMLM modalities: PALM, dSTORM and PAINT. For later analysis, nano-org automatically assesses the localisation density and coverage. Nano-org then allows users to navigate through the publicly accessible datasets using metadata tags and download pertinent data for their own analysis (Figure 1).

**Figure 1:**
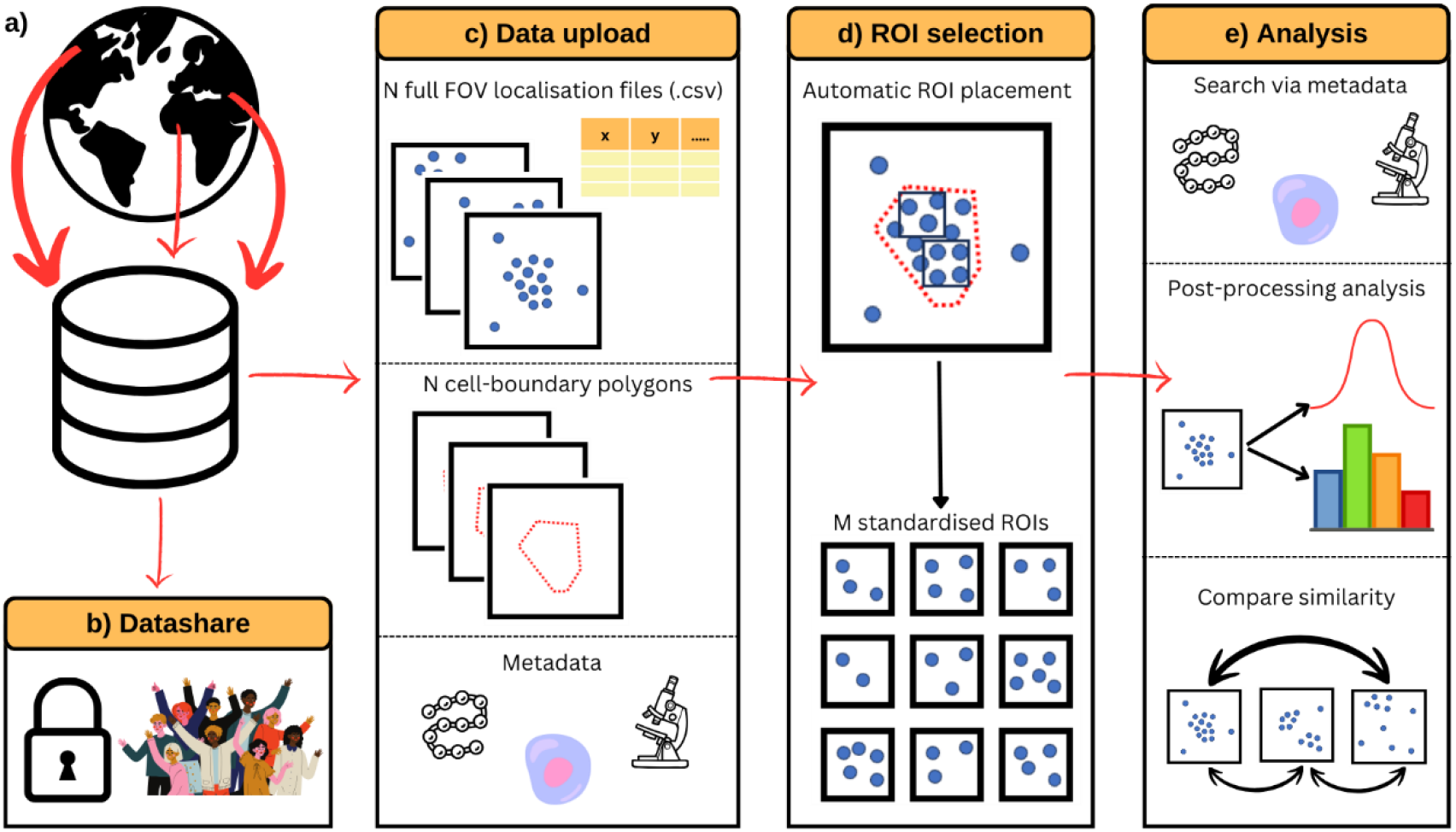
Key functionalities of nano-org. **a)** Users can upload their data, **b)** selecting from various privacy settings. **c)** Users can upload localisation files (.csv) along with cell-bounding polygons. Metadata requirements ensure comprehensive dataset documentation and enhance search functionality. **d)** Uploaded data is split into regions of interest (ROIs) for downstream analysis. **e)** The database enables users to explore public datasets, extract relevant information, and utilise statistical similarity tools for comparative analysis.

A key feature of nano-org is the ability to search its contents by the statistical similarity of datasets (Figure 2). This means users can upload a condition and search for other conditions where proteins exhibit the most similar nanoscale organisation. This is analogous to searching a gene sequence database based on sequence homology. Briefly, on upload, ROIs are subdivided into 30nm^2^ bins (optimised and chosen based on the average precision of localisations). We then form a frequency histogram for the number of localisations in each bin and construct the empirical cumulative distribution function (CDF) of the set of frequencies (histogram heights). Every 5 minutes, a check is run which first identifies all new or modified data in the database. Then, for every pair of ROIs, we compute the largest discrepancy between their CDFs, as in the Kolmogorov-Smirnov (K-S) test [9, 10]. The K-S test yields a dissimilarity value (λ), where an increasing λ denotes greater dissimilarity between the ROIs. For condition-wide comparisons the mean dissimilarity, 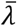, is computed. This procedure results in a list of all other database contents ranked by their nanoscale organisational similarity to the condition in question. This list can then be downloaded and further filtered or searched using metadata. Two different versions of this ranked list are produced – one as described above and one which aims to be invariant to differences in the total number of localisations in the ROI (e.g. if the user has not controlled for expression level, acquisition time etc). This is achieved by thinning uploaded datasets to a standard value (30 localisations/*μ*m^2^); the minimum density at which the algorithm could identify meaningful differences between datasets while also minimising the exclusion of sparse datasets. The system scales efficiently with the number of ROIs in the database, as it compares pre-computed histograms rather than raw data, enabling many comparisons within a reasonable timeframe. The infrastructure supports real-time updates and comparisons, ensuring timely results even as the database grows. A complete description of the algorithm is provided in the Supplementary Material.

**Figure 2:**
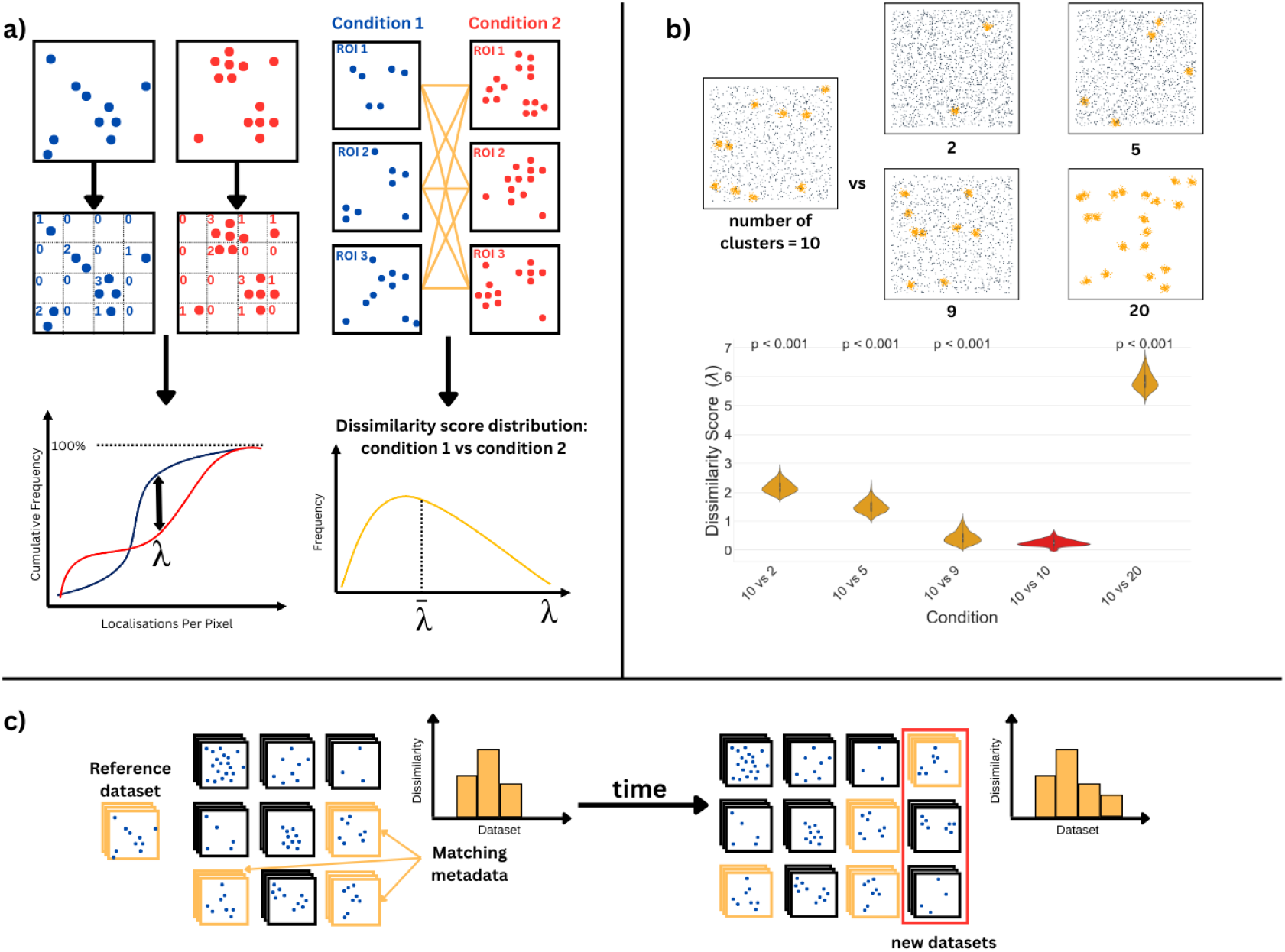
Nano-org’s similarity search approach. **a)** Two ROIs are divided into 30nm^2^ bins to generate cumulative frequency histograms of localisations. A K-S test yields a dissimilarity value (λ) between the ROIs, with mean dissimilarity 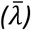 calculated across datasets for comparison. **b)** Example of simulated data showing increasing dissimilarity as the number of clusters varies from 10, while keeping the total localisations per ROI constant. **c)** Real-time analysis on nano-org enables continuous comparison with public datasets, filterable by metadata. Ranked similarity lists are updated with new uploads.

Our similarity scoring method underwent rigorous testing on simulated data. For example, ROIs with 10 clusters were compared to ROIs with different numbers of clusters while keeping the total number of localisations the same. Dissimilarity increased with the difference in the number of clusters (Figure 2) and statistical testing demonstrated that the differences were statistically significant. Testing with different cluster sizes and different numbers of points per cluster was well as with simulated fibrous data mimicking the nanoscale organisation of the cytoskeleton is shown in Supplementary Figures S1 and S2.

To illustrate the utility of our approach, we investigated the organisation of one of the datasets stored on the database - T cell immunoreceptor with immunoglobulin and ITIM domains (TIGIT); imaged using dSTORM. TIGIT is an inhibitory receptor on various immune cells, including T and NK cells. Recent findings show that upon ligation, TIGIT forms nanoclusters co-localised with the activating T cell receptor, and this clustering is important for its signal transduction [11]. From the ranked similarity list, we found that TIGIT organisation in NK cells was more similar to TIGIT in other cell types, specifically Jurkat and CD4+ T cells than it is to other proteins, such as KIR2DL1 and NKp30, on the surface of NK cells (Figure 3). In fact, dissimilarity values comparing TIGIT in NK cells to TIGIT in Jurkat cells were not statistically significant. This suggests that the protein identity, rather than cell type, is most important in defining the nanoscale organisation in this case. Rankings are preserved after thinning the data showing the trends are due to genuine differences in the protein nanoscale distribution and not solely due to differences in expression levels.

**Figure 3:**
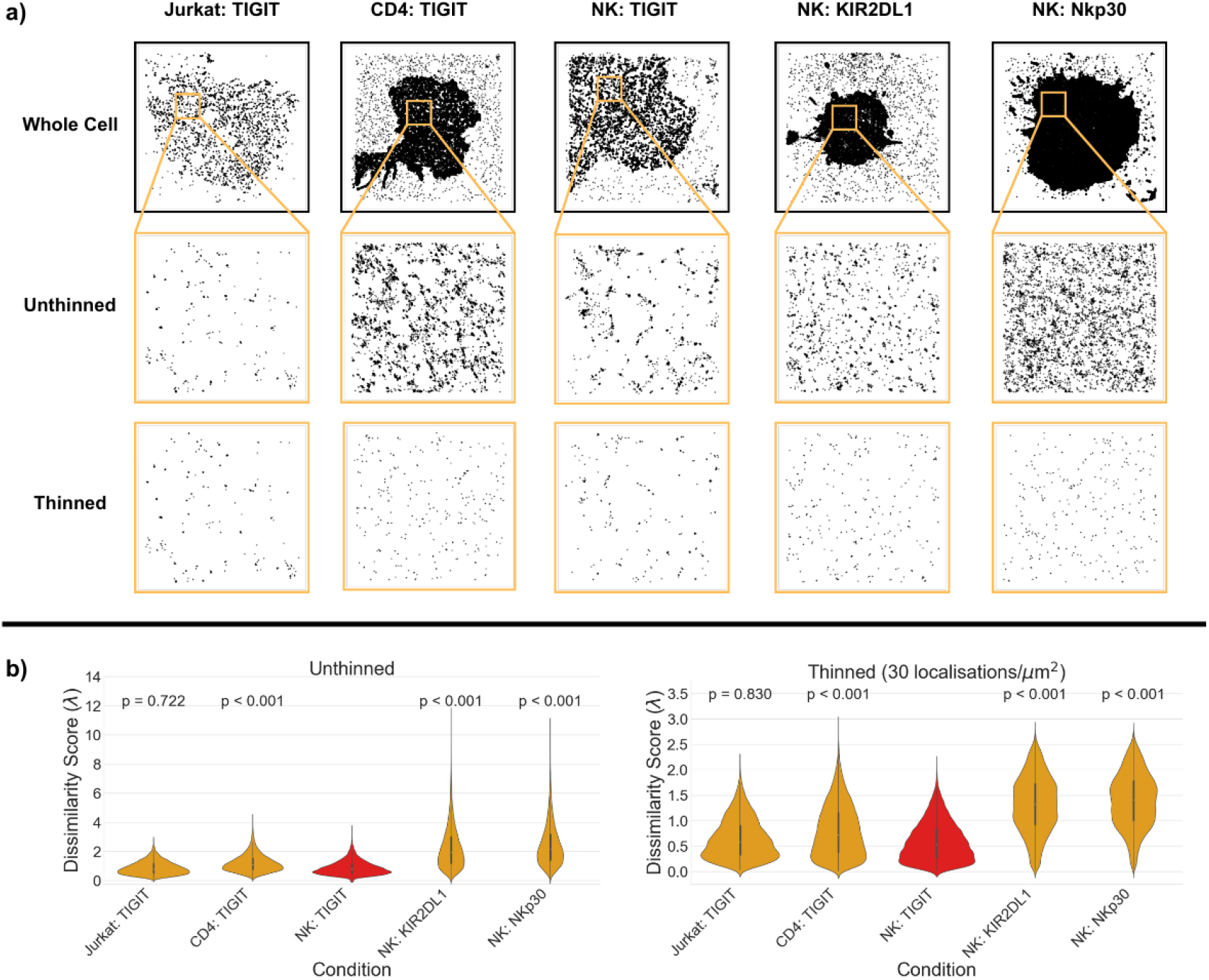
Dissimilarity scores between experimental data in nano-org. **a)** Examples of whole cell coordinates used for analysis, along with example ROIs for each condition of interest: TIGIT in Jurkat cells, CD4 cells, and NK cells; NKp30 in NK cells;, and KIR2DL1 in NK cells. These ROIs are presented either unthinned or thinned to 30 localisations/μm^2^. **b)** Dissimilarity between TIGIT in NK cells with itself (red) and with all other conditions.

In conclusion, nano-org is a publicly accessible and curated database of SMLM data, designed to facilitate collaborative data sharing, enhancing accessibility and reproducibility. Its unique framework allows searches based on statistical similarity, enabling investigations into the biophysical mechanisms of nanoscale organisation and the effects of mutations or treatments on protein distributions. This resource represents the first step in developing spatial nano-omics – the systematic study of cellular nanoscale architecture.

## Acknowledgements

The research described in this paper was carried out with the assistance of Advanced Research Computing at the University of Birmingham. This included support from the Research Software Group to develop the database and website, data storage on the Research Data Store, computations on the BlueBEAR HPC service and use of a BEAR Cloud Virtual Machine to host the website. DMO and SS acknowledge funding from the Biotechnology and Biological Sciences research Council (BBSRC) grant BB/X018644/1. RH received funding from the European Research Council (ERC) through grant 101001332-SelfDriving4DSR and Horizon Europe through grants 101057970-AI4LIFE and 101099654-RTSuperES. RH also acknowledge the support of the Gulbenkian Foundation (Fundação Calouste Gulbenkian). Views and opinions expressed are, however, those of the authors only and do not necessarily reflect those of the European Union. Neither the European Union nor the granting authority can be held responsible for them. This work was also supported by a European Molecular Biology Organization (EMBO) installation grant (EMBO-2020-IG-4734 to RH).

## Author Contributions

SS, DJN, MHHB, KS and JAP developed and tested the algorithms. JAP, MAA and AG implemented the database and website. DJN, JW, KSH, DJW, RP and DMD provided simulations, experimental data and testing. PRD supported with statistical analysis. RH, SFL and DMO conceived the work. SS and DMO wrote the manuscript.

## Code and data availability

All experimental data is stored and available for download on https://nano-org.bham.ac.uk. The implementation of the website and database is available at https://gitlab.bham.ac.uk/owendz-protein-databank/nano-org-website. The core analysis functionality and algorithms used by nano-org are implemented as a stand-alone python package which is available at https://gitlab.bham.ac.uk/owendz-protein-databank/smlm-analysis. All Python scripts used to produce simulated data and violin plots in figures and supplementary figures are available at https://gitlab.bham.ac.uk/shirgils/nano-org-similarity-scoring.

