## Supplemental Information for "Nano-org, a functional resource for single-molecule localisation microscopy data"

**Supplementary material**

### Nano-org: Technical Description

Nano-org is a Django-based website, with an SQLite database backend, which is hosted on a BEARCloud virtual machine at the University of Birmingham. Celery and RabbitMQ are used to schedule background tasks including checking the database for new and modified data and initiating computationally intense analysis tasks via job submission to BlueBEAR - the University of Birmingham's supercomputer for high performance computing (HPC). Celery tasks are also used to retrieve analysis results from HPC jobs and incorporate them into the website and database. Uploaded data and analysis results are stored on the University of Birmingham's central Research Data Store and made available to download through the website. Core analysis functionality, including cumulative histogram generation and KS score calculation, is incorporated into our stand-alone python package smlm. This package is utilised by nano-org but can also be used by researchers who want to develop their own customised analysis pipelines.

### Dissimilarity algorithm

To compare the dissimilarity between two ROIs, let $F(x)$ and $G(x)$ be their empirical cumulative distributions with sample sizes $m$ and $n$ respectively. Here, the sample sizes are the number of 30x30 nm bins within the ROI that contain at least one localisation. A two-sample Kolmogorov-Smirnov (K-S) Statistic is employed with the following modifications:

Using the definition,

$D_{mn} > c(\alpha)\sqrt{J}$ , where,

$$D_{mn} = \max_{x} |F(x) - G(x)|,$$

$$J = \frac{n+m}{nm},$$

$$c(\alpha) = \sqrt{-\frac{1}{2}\ln(\frac{\alpha}{2})} .$$

Here, $\alpha$ is the confidence level, where we set $\alpha = 0.05$. We define,

$$KS = \frac{D_{mn}}{c(\alpha)\sqrt{J}} = \lambda$$

Where $\lambda$ is then the dissimilarity score between the two datasets.

### Summary of Statistical Testing Method

We obtain a p-value for the set of dissimilarities using a permutation test. First, we compute the test statistic, T, as the mean inter-group dissimilarity minus the two mean intra-group dissimilarities. Next, under N = 1000 simulations, we randomly re-assign the ROIs into two groups of the same size, and recompute the test statistic, T*. We return the proportion of simulations for which T* >= T.

**Supplementary figures**

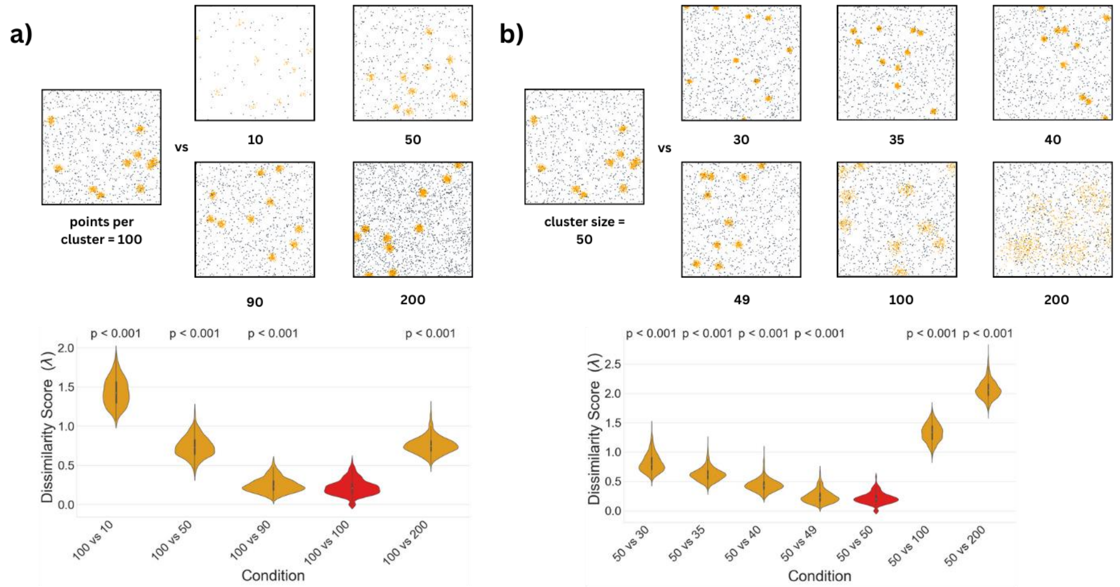


**S1: Dissimilarity scores between simulated Gaussian clusters**. **a)** Example ROIs for each Gaussian cluster simulations with different numbers of points per cluster with dissimilarity scores comparing 100 points per cluster with all other conditions. **b)** Example ROIs for each Gaussian cluster simulations with different cluster sizes with dissimilarity scores comparing clusters of size 50 nm with itself and all other conditions.

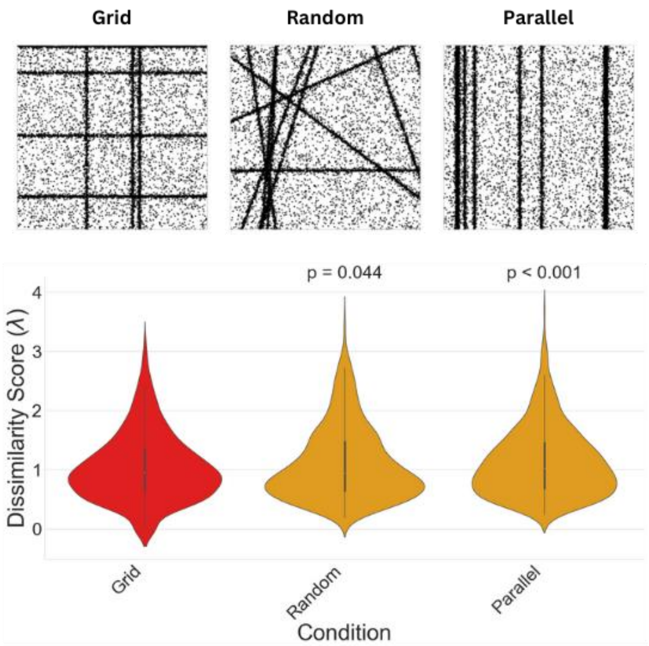


**S2: Dissimilarity scores between simulated fibres**. Fibres are arranged on a grid, randomly, or parallel to each other. Dissimilarity scores between fibres arranged on a grid are compared to themselves and with randomly oriented and parallel fibres.
